# Effects of Neighboring Phosphorylation Events on the Affinities of pT181-Tau Antibodies

**DOI:** 10.1101/2023.09.10.557075

**Authors:** Sehong Min, Rodrigo Mohallem, Uma K Aryal, Tamara L. Kinzer-Ursem, Jean-Christophe Rochet

## Abstract

A tau variant phosphorylated on threonine 181 (pT181-tau) has been widely investigated as a potential Alzheimer’s disease (AD) biomarker in cerebrospinal fluid (CSF) and blood. pT181-tau is present in neurofibrillary tangles (NFTs) of AD brains, and CSF levels of pT181-tau correlate with overall NFT burden. Various immuno-based analytical methods, including Western blotting and ELISA, have been used to quantify pT181-tau in human biofluids. The reliability of these methods depends on the affinity and binding specificity of the antibodies used to measure pT181-tau levels. Although both of these properties could in principle be affected by phosphorylation within or near the antibody’s cognate antigen, such effects have not been extensively studied. Here, we developed a biolayer interferometry assay to determine the degree to which the affinity of pT181-tau antibodies is altered by the phosphorylation of serine or threonine residues near the target epitope. Our results revealed that phosphorylation near T181 negatively affected the binding of pT181-tau antibodies to their cognate antigen to varying degrees. In particular, two of three antibodies tested showed a complete loss of affinity for the pT181 target when S184 or S185 was phosphorylated. These findings highlight the importance of selecting antibodies that have been thoroughly characterized in terms of affinity and binding specificity, addressing the potential disruptive effects of post-translational modifications in the epitope region, to ensure accurate biomarker quantitation.

---

Alzheimer’s disease (AD) is a neurodegenerative disorder clinically characterized by memory loss and cognitive decline.^1, 2^ Neuropathological hallmarks of AD include extra-cellular plaques and intracellular neurofibrillary tangles, consisting of fibrillar deposits of the amyloid-beta peptide and hyperphosphorylated forms of the microtubule-binding protein tau, respectively.^3^ AD diagnosis relies largely on the evaluation of measures of cognitive impairment.^4^ However, plaques and tangles affecting the brains of AD patients are thought to begin developing decades before clinical symptoms appear, and this underlying pathology cannot be reversed with currently available treatments.^3, 4^ These observations imply that early diagnosis and intervention to delay the onset of AD and prevent further disease progression would be highly beneficial.

A major priority in the AD field is to identify peripheral biomarkers that report on early stages of the disease. According to the NIA-AA (National Institute on Aging -Alzheimer’s Association) research framework, AD can be detected by assaying biomarkers indicative of neuropathologic changes, including amyloid-beta (A), pathological tau (T), and neurodegeneration (N), collectively referred to as the ATN classification system.^5^ Early stages of AD may be characterized by increases in the levels of these biomarkers in cerebrospinal fluid (CSF) or blood prior to the onset of symptoms. In particular, assays of amyloid-beta and tau in CSF are currently used to aid in the diagnosis of AD.^6^ Moreover, recent findings suggest that blood levels of amyloid-beta and tau are reliably associated with AD progression and can be used to predict neuropathologic changes in the brain before symptoms develop.^7, 8^

Elevated levels of phosphorylated tau (p-tau) observed in the CSF and blood of AD patients reflect the presence of tau pathology in the brain.^7, 9^ Although pathological tau in the brains of AD patients is phosphorylated at multiple sites,^10^ most p-tau assays are designed to quantify tau phosphorylated at T181 because of its strong correlation with cognitive impairment, amyloid-beta accumulation, and tau pathology detected by positron emission tomography (PET).^11-14^ pT181-tau in CSF is widely considered a promising biomarker to facilitate AD diagnosis and the monitoring of disease progression,^6, 13, 15^ and evidence suggests that blood pT181-tau can also serve as a biomarker to predict tau and amyloid-beta pathologies and differentiate AD from other disorders.^7^

Measurements of pT181-tau levels in CSF and blood rely heavily on immunoassays including ELISA,^14, 16^ single molecule array (SiMoA),^17, 18^ electrochemiluminescence immunoassay (ECLIA),^19^ and xMAP technology.^20^ Surface plasmon resonance (SPR) has also been used as an antibody-based method for tau quantification.^21, 22^ These immunoassays depend critically on the ability of tau antibodies to recognize their cognate antigens in biofluids with high sensitivity, selectivity, and specificity. Immunoassays for pT181-tau quantification have been optimized primarily with respect to antibody sensitivity (with the aim of enabling the measurement of femtomolar levels of p-tau in biological samples) rather than antibody specificity.^23^

Mass spectrometry (MS) analyses have revealed that residues near pT181-tau are phosphorylated in AD.^24-26^ Tau was found to be phosphorylated on both T175 and T181 in the brains and CSF of AD patients.^24-26^ The protein has also been shown to be phosphorylated on residues S184 and S185 in AD brain, albeit at a low frequency (<5%). Despite this evidence that tau variants associated with AD can be phosphorylated on residues near pT181, the effects of these neighboring phosphorylation events on the affinity of pT181-tau antibodies for their cognate antigen have not been investigated.

In this study, we used a novel biolayer-interferometry (BLI) assay to determine the effects of phosphorylation events near residue pT181 on the affinities and binding specificities of three pT181-tau antibodies. The BLI data revealed that phosphorylation near T181 reduced the binding of pT181-tau antibodies to their target to varying degrees. These findings emphasize the importance of selecting antibodies that have been rigorously assessed for potential disruptive effects of post-translational modifications near the target antigen to avoid pitfalls associated with false-positive or false-negative readings in tau biomarker assays. Moreover, the BLI method described here can serve as a powerful platform for the quantitative assessment of tau antibody affinity and specificity.

## EXPERIMENTAL SECTION

### Materials

All chemicals were purchased from Sigma-Aldrich (St. Louis, MO) unless otherwise stated. Octet® SAX high precision streptavidin biosensors and Octet® 384TW tilted-bottom plates were purchased from Sartorius (Göttingen, Germany). Peptides were synthesized by GenScript (Piscataway, NJ). Trypsin and peptide desalting columns were obtained as Pierce products from Thermo Fisher Scientific (Waltham, MA). The pRK172-h-tau-2N4R bacterial expression plasmid was a gift from David Eliezer (Weill Cornell Medical College).

### Antibodies

The following antibodies specific for pT181-tau were used in this study: (i) mouse monoclonal AT270, purchased from Thermo Fisher Scientific; (ii) rabbit monoclonal D9F4G, purchased from Cell Signaling Technology (CST, Danvers, MA); and (iii) rabbit monoclonal E4G7X, provided by Thorsten Wiederhold and Arica Aiello (CST). The following antibodies were used for Western Blot analysis in addition to the anti-pT181-tau antibodies: (i) chicken anti-tau (#1998-TAU), obtained from PhosphoSolutions (Aurora, CO), (ii) goat anti-mouse IgG IRDye 680RD, (iii) donkey anti-rabbit IgG IRDye 680RD, and (iv) donkey anti-chicken IgG IRDye 800CW, obtained from LI-COR Biosciences (Lincoln, NE).

### Biolayer Interferometry (BLI)

Prior to carrying out the BLI measurements, the SAX biosensors were hydrated in a 96-well plate by incubating in BLI assay buffer, consisting of PBS (10 mM phosphate buffer, 2.7 mM KCl, and 137 mM NaCl, pH 7.4) with 0.1% (w/v) BSA and 0.05% (v/v) tween-20 (200 μL per well), for 10 min at 22°C. Tau peptide was biotinylated at the N-terminus with an aminohexanoic acid linker (Figure 1A). Peptide solutions used for the loading step were prepared by reconstituting lyophilized tau peptides in H_2_O at a concentration of 1 mg/mL and diluting to a final concentration of 10 ng/mL in assay buffer. BLI measurements were performed by loading 40 μL samples to the wells of a 384-well tilted bottom plate and shaking the plate at 1,000 rpm in the Octet RED384 instrument (ForteBio) at 30°C using the protocol shown in Figures 1B and S1. To obtain reliable kinetic constants, a dilution series of seven antibody concentrations as well as a reference well containing only assay buffer in the absence of antibody were run in parallel in the association phase. The BLI data were analyzed and fit to kinetic curves using the Octet Data Analysis software. All BLI experiments were performed at least in duplicate. Both peptides and antibodies were freshly diluted in assay buffer before each experiment.

**Figure 1.**
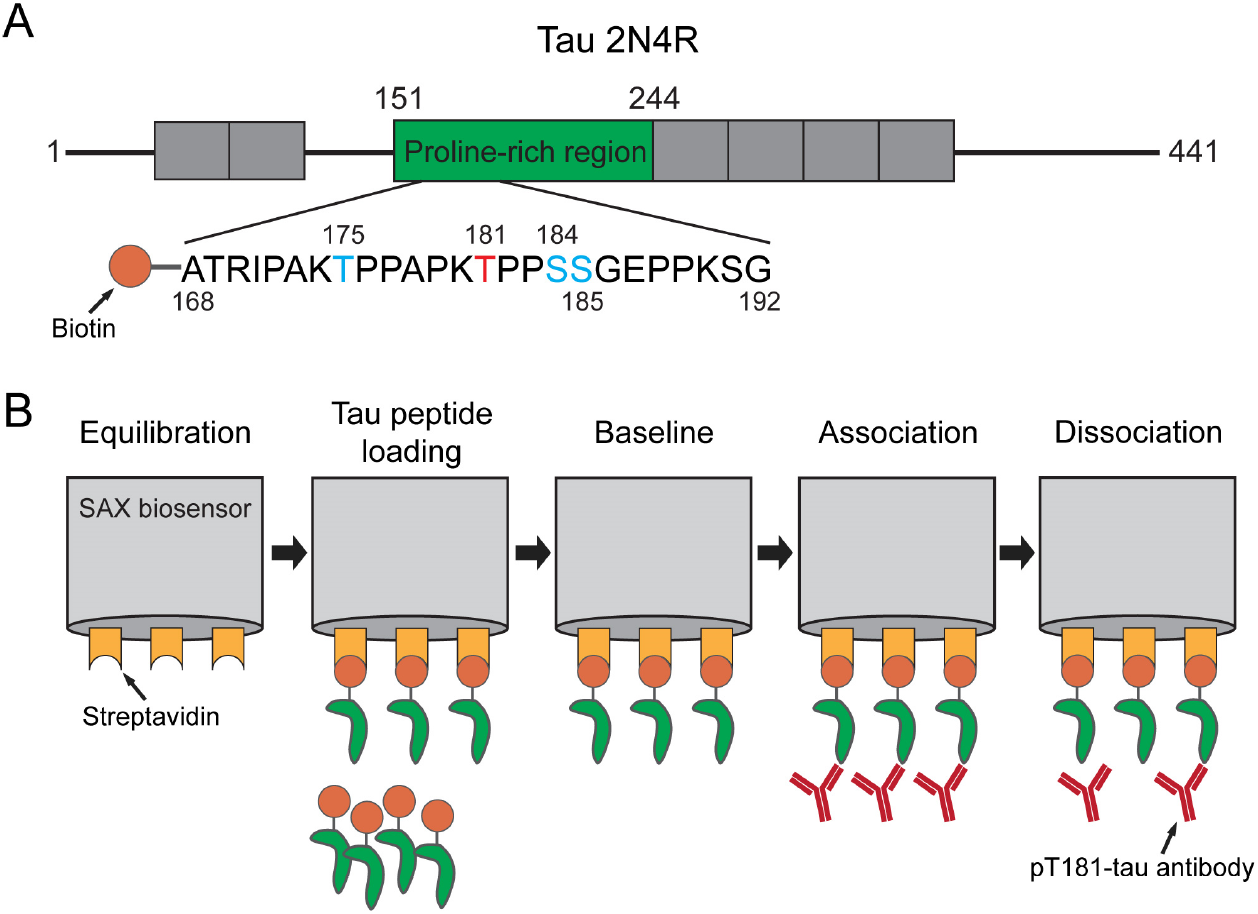
Design of biolayer interferometry (BLI) assay to determine the affinities and binding specificities of pT181-tau antibodies for tau phosphopeptides. (A) Schematic representation of N-terminally biotinylated tau peptides used as targets peptides in the BLI assay. The peptides span residues 168 to 192 and are phosphorylated on T181 as well as different combinations of T175, S184, and S185. (B) Schematic representation of the steps involved in the BLI assay. Streptavidin-coated (SAX) biosensors are equilibrated in wells containing the BLI assay buffer (Equilibration) and then transferred into wells containing assay buffer plus biotinylated tau target peptide to immobilize the peptide on the biosensor (Tau peptide loading). Peptide-loaded biosensors are transferred into wells containing assay buffer for washing and to measure the baseline signal (Baseline). The biosensors are then dipped into wells containing assay buffer plus pT181-tau antibody (Association), before being transferred back into wells containing BLI assay buffer without antibody (Dissociation).

### Expression and Purification of Non-Phosphorylated Tau

Non-phosphorylated, human tau 2N4R was generated using an *E. coli* expression system. Briefly, *E. coli* BL21(DE3) cells transformed with pRK172-h-tau-2N4R were incubated in LB media supplemented with ampicillin (100 μg/mL) and IPTG (1 mM) for 4 h at 37°C to induce protein expression. The cells were pelleted by centrifugation at 6,000 *g* for 15 min at 4°C, resuspended in lysis buffer (20 mM MES, 0.2 mM MgCl_2_, 1 mM EGTA, 0.25 mg/mL lysozyme, 1 μg/mL DNase I, pH 6.8), and lysed with a French press cell disruptor at 4°C. After boiling the lysate for 20 min, denatured proteins were pelleted by centrifugation at 30,000 *g* for 30 min at 4°C, and the supernatant was loaded onto a HiPrep SP HP column on an AKTA pure chromatography system. The column was equilibrated with cation exchange buffer (20 mM MES, 50 mM NaCl, 1 mM MgCl_2_, 1 mM EGTA, 2 mM DTT, 0.1 mM PMSF, pH 6.8), and proteins were eluted with a linear gradient ranging from 50 mM to 1 M NaCl. The resulting protein solution was purified further using a HiLoad 16/600 Superdex 200 pg column with 1.5 column volumes of PBS (10 mM phosphate buffer, 2.7 mM KCl, and 137 mM NaCl, pH 7.4), and fractions containing tau (identified by SDS-PAGE with Coomassie blue staining) were pooled and stored at -80°C.

### Expression and Purification of Phosphorylated Tau

Phosphorylated tau was produced using a baculovirus/Sf9 insect cell system. A cDNA encoding human tau 2N4R (excised from pRK172-h-tau-2N4R) was inserted into the baculovirus transfer vector pVL1393. Sf9 insect cells were cotransfected with each of the pVL-2N4R constructs and BestBac DNA using the BestBac 2.0 Δ v-cath/chiA Baculovirus Cotransfection Kit to generate baculoviruses encoding the different tau-2N4R variants. Sf9 cells in suspension culture were transduced with tau-2N4R baculovirus at a multiplicity of infection (MOI) of 5, incubated for 72 h at 27°C, and harvested by centrifugation. The pelleted cells were resuspended in 10 volumes (v/w) of lysis buffer (50 mM Tris-HCl, 500 mM NaCl, 5 mM DTT, 10 mM EGTA, 20 mM NaF, 1 mM Na_3_VO_4_, 5 μM microcystin, protease inhibitor cocktail (P8340, Sigma-Aldrich), pH 7.4) and lysed in a Dounce homogenizer with 20 strokes. The lysate was supplemented with 1% (w/v) streptomycin sulfate, incubated for 30 min at 4°C, and cleared by centrifugation to pellet the insoluble genomic DNA (30,000 *g* for 30 min at 4°C). The supernatant was then boiled for 10 min, cleared by centrifugation to pellet denatured proteins (30,000 *g* for 30 min at 4°C), and dialyzed overnight against anion exchange buffer (100 mM MES, 2 mM DTT, 1 mM EGTA, 1 mM MgSO_4_, 0.1 mM PMSF, pH 6.8). The dialysate was loaded onto a HiPrep Q HP column, and proteins were eluted with a linear gradient ranging from 50 mM to 1 M NaCl. Fractions containing tau were processed as described above for the non-phosphorylated protein. Phosphomimetic tau variants were produced by GenScript.

### Western Blot Analysis

Western blot analysis of tau-2N4R variants was carried out by mixing solutions of recombinant protein in a 1:1 (v/v) ratio with 2x Laemmli sample buffer, adding β-mercaptoethanol to a final concentration of 5% (v/v), and boiling the resulting mixtures for 5 min. Proteins were separated via SDS-PAGE through a 12% (w/v) Tris-glycine polyacrylamide gel and transferred to an Immobilon^®^-FL PVDF membrane. After blocking in Intercept^®^ (TBS) blocking buffer, the membrane was incubated overnight at 4°C with anti-tau (1:2,000) and one of anti-pT181-tau (1:1,500). After washing in TBS (25 mM Tris buffer, 137 mM NaCl, pH 7.6) plus 0.1% (w/v) Tween 20 (TBS-T) three times, the membrane was probed for 1 h at 22°C with the IRDye 800CW secondary antibody donkey anti-chicken IgG (1:10,000) for blots probed with anti-tau and the IRDye 680RD secondary antibody goat anti-mouse IgG (1:10,000) for blots probed with AT270, or the IRDye 680RD secondary antibody donkey anti-rabbit IgG (1:10,000) for blots probed with D9F4G or E4G7X. After washing with TBS-T three times, the membrane was scanned using an Odyssey M LI-COR scanner, and band intensities were quantified using ImageJ.

### Statistical Analysis

Statistical analyses were conducted using GraphPad Prism 9 (La Jolla, CA). All values are expressed as the mean ± SEM.

## RESULTS AND DISCUSSION

### Development of a BLI Assay to Measure pT181-Tau Antibody Affinities

Our first objective was to develop and optimize a BLI assay that could be used to reliably measure the affinities of pT181-tau antibodies, including the three antibodies examined here: AT270, D9F4G, and E4G7X. The assay (Figure 1) involves (i) immobilizing N-terminally biotinylated tau peptides spanning amino acid residues 168-192 (based on the numbering of the longest human tau isoform 2N4R; Figure 1A) onto streptavidin-coated biosensors; (ii) establishing a stable baseline signal; (iii) monitoring the binding of pT181-tau antibody to the immobilized peptides at different antibody concentrations (association phase); and (iv) monitoring the release of antibody from the peptide in buffer minus antibody (dissociation phase). The instrument provides a real-time measure of the shift in optical interference (reported as a shift in wavelength, in nm) resulting from the binding of peptide or antibody to the biosensor tip. To analyze the data, we subtracted the reference signal (measured in the absence of antibody) from the raw data values obtained with each biosensor, and the data were fitted to a 1:1 binding model. Local fitting was first performed to compute the kinetic constants for the curve obtained at each antibody concentration (i.e., k_on_ and k_off_ values relating to the association and dissociation phases, respectively, from which the equilibrium dissociation constant (K_D_) was determined as the ratio of k_off_ to k_on_). Kinetic traces with measurable k_on_ and k_off_ values, R^2^ > 0.95, and χ^2^< 3 were selected for a subsequent global fit analysis to obtain overall k_on_, k_off_, and K_D_ values spanning multiple antibody concentrations.

In developing our BLI assay, we first assessed the optimal concentration of tau peptide loaded onto the streptavidin-coated biosensors. To measure an antibody’s affinity for its cognate antigen accurately via BLI, it is critical to ensure minimal dissociation of the peptide from the biosensor to which it is bound.^29^ Overloading of the peptide can lead to an unstable baseline that reflects ongoing peptide dissociation, resulting in a poor fit of the antibody binding and dissociation data to a 1:1 model. The pT181-tau peptide was added to the biosensors at concentrations of 10 to 1000 ng/mL, and the AT270 antibody was then added at 1.5 nM during the association phase (Figure S2A). As all peptide concentrations except 10 ng/mL showed unstable baselines, for the next trial, the biosensors were incubated with pT181-tau peptide at 1 to 10 ng/mL during the loading phase, followed by the AT270 antibody at 1, 1.5, or 2 nM during the association phase (Figure S2B). Peptide loading at 10 ng/mL produced a wavelength shift of ∼0.1 nm during the loading phase without any evidence of drift or peptide detachment from the biosensor, and a wavelength shift of ∼0.5 to 1 nm during the association phase, indicating that the peptide was properly immobilized for subsequent antibody binding. Accordingly, this peptide loading concentration was used in all of the remaining analyses.

Next, we determined the optimal range of concentrations of pT181-tau antibody to be added during the BLI association phase. Here, our goal was to identify antibody concentrations that produced a maximum signal occurring between the lower limit of detection and the saturation signal, based on the assumption that kinetic constants obtained by fitting curves within this range would be most accurate.^30^ The optimal antibody concentration range was found to be 100 to 2,000 pM for AT270, 200 to 4,000 pM for D9F4G, and 400 to 4,000 pM for E4G7X (Figure S3). These concentration ranges were used for all of the remaining analyses.

Having optimized our BLI assay with respect to the peptide and antibody concentrations, we then assessed the specificity of each of the three antibodies for pT181-tau peptide versus the corresponding unphosphorylated peptide target. All three pT181-tau antibodies bound to the phosphorylated peptide in a concentration-dependent manner, resulting in an increase in the maximum wavelength shift evident in the sensorgram traces with increasing antibody concentration (Figures 2A and S3). In contrast, none of the three antibodies exhibited detectable binding to the unphosphorylated tau peptide (Figure S4). The order of antibody binding affinities for the pT181-tau peptide target was AT270 > D9F4G > E4G7X, with estimated K_D_ values of 63 ± 13 pM, 360 ± 35 pM, and 459 ± 37 pM, respectively (mean ± SEM; Figure 2B and Table S1). The higher affinity of AT270 was the result of a higher k_on_ value (Figure 2C) and a lower k_off_ value (Figure 2D). These results indicate that (i) our BLI assay can be used to measure differences in the binding affinities of pT181-tau antibodies with K_D_ values spanning ∼60 to ∼500 pM; and (ii) all three of the pT181-tau antibodies tested here bound to the phosphorylated peptide target with high specificity but a range of affinities, as reflected by their K_D_ values that varied over an ∼8-fold range.

**Figure 2.**
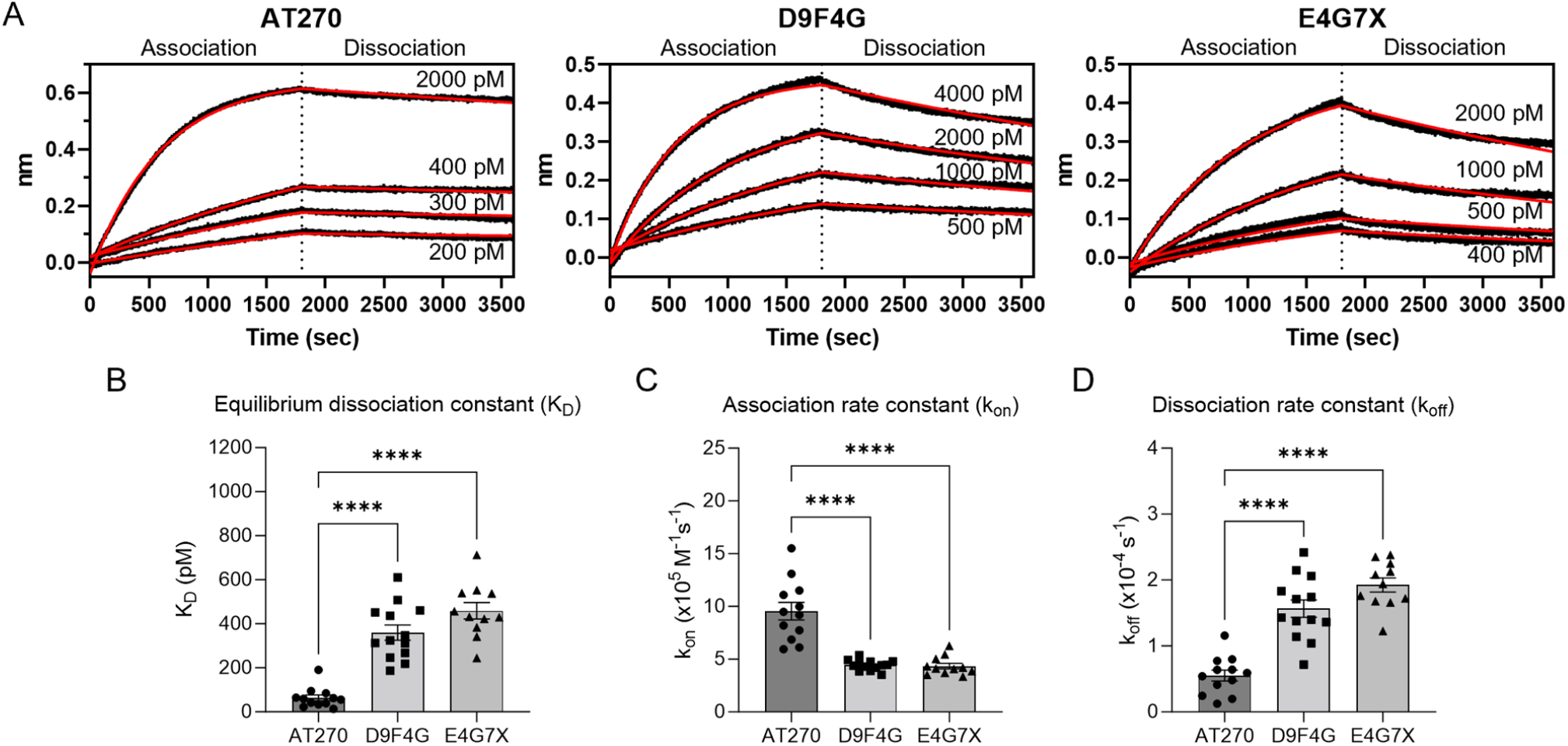
Measurement of antibody affinities for the pT181-tau peptide. (A) Sensorgrams showing raw-data traces (black) and best-fit curves (red) corresponding to the association and dissociation of the pT181-tau antibody AT270 (left), D9F4G (middle), or E4G7X (right) upon incubation at different concentrations with biosensors loaded with pT181-tau peptide. The best-fit curves were obtained using a 1:1 model with a global-fit analysis. (B-D) Bar graphs showing equilibrium dissociation constants (K_D_ values) (B), association rate constants (k_on_ values) (C), and dissociation rate constants (k_off_ values) (D) determined for AT270, D9F4G, and E4G7X. The lower K_D_ value observed for AT270 is a reflection of this antibody’s higher k_on_ value and lower k_off_ value. All values are expressed as the mean ± SEM (n = 11-13); ^****^p ≤ 0.0001, one-way ANOVA followed by Tukey’s multiple comparisons post hoc test. Each data point represents the value obtained from one independent, replicate experiment.

### Effects of Neighboring Phosphorylation Events on pT181-Tau Antibody Affinities

In the next phase of our study, we examined how the affinity of each pT181-tau antibody for its peptide target was affected by phosphorylation events near the pT181 epitope (referred to here as ‘neighboring’ phosphorylation events). To address this question, we monitored the binding of the pT181-tau antibodies to a panel of tau peptides phosphorylated on T181 as well as different combinations of neighboring residues (T175, S184, and/or S185; Figure 3 and Table S1) using the BLI assay. In one set of analyses, we assessed the ability of the pT181-tau antibodies to bind to pT181-tau peptides that were doubly phosphorylated (pT175/pT181, pT181/pS184, and pT181/pS185) or triply phosphorylated (pT175/pT181/pS184, pT175/pT181/pS185, and pT181/pS184/pS185). The results revealed that neighboring phosphorylation events had no effect on AT270 binding affinity, whereas D9F4G and E4G7X antibodies failed to bind to the pT181-tau peptides when either S184 or S185 was phosphorylated. Taken together, these data show that the binding of pT181-tau antibodies to the peptide target is disrupted to different extents by neighboring phosphorylation events.

**Figure 3.**
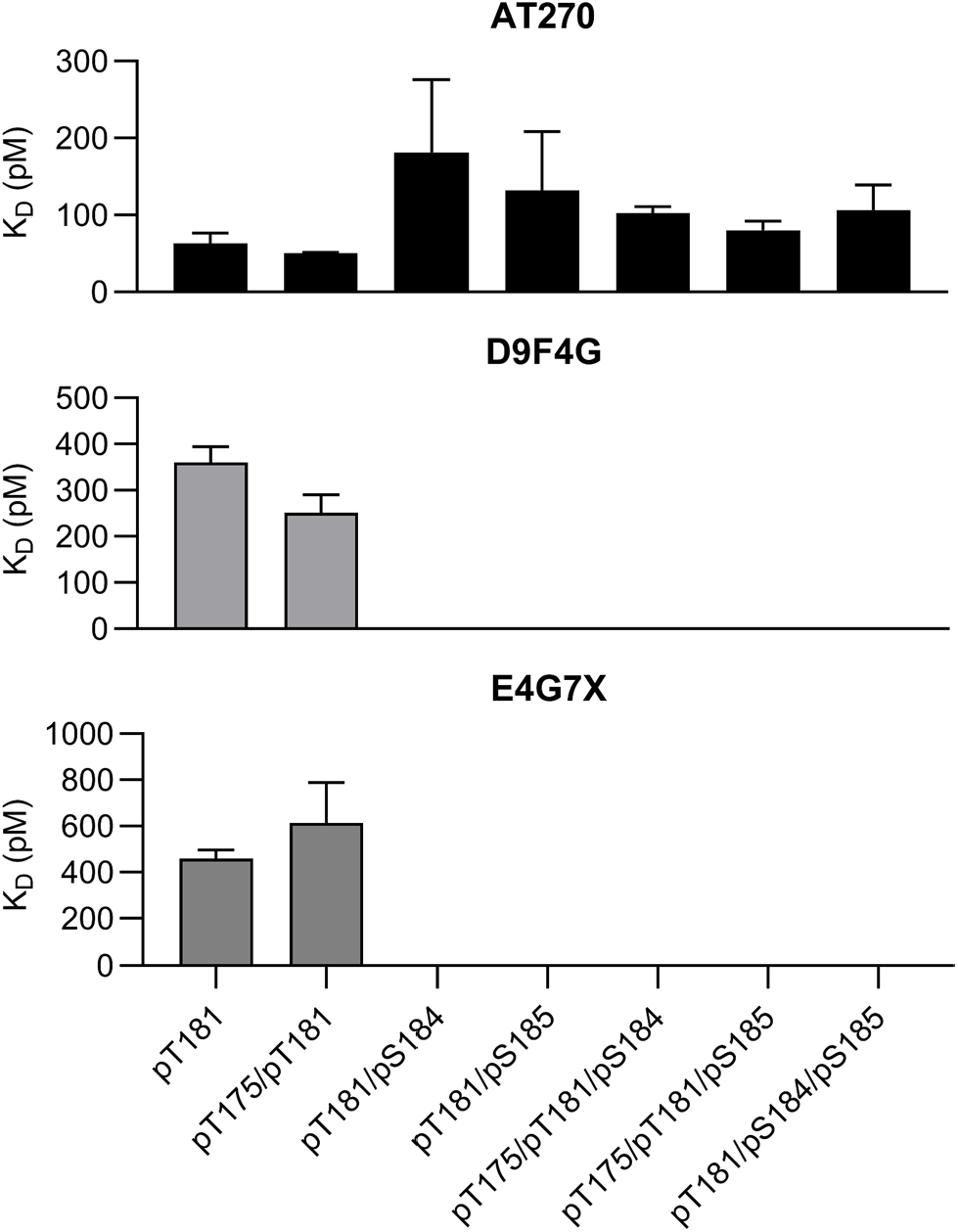
The affinities of pT181-tau antibodies for the antigenic peptide are affected to different extents by neighboring phosphorylation events. Bar graph showing equilibrium dissociation constants (K_D_ values) determined for AT270, D9F4G, and E4G7X with respect to multiply-phosphorylated tau peptides. The binding of D9F4G and E4G7X to pT181-tau peptide was more severely disrupted by neighboring phosphorylation events than that of AT270. Values are expressed as the mean ± SEM (n = 2-13) and are also listed in Table S1. The absence of a bar indicates that no measurable antibody binding was detected. The data obtained for peptide pT181 are the same as those shown in Fig. 2.

Based on evidence that some pathological forms of tau in biofluids or the postmortem brains of AD patients are phosphorylated at S184 or S185, albeit at a low abundance,^24^ our findings suggest that assays involving D9F4G or E4G7X could potentially lead to underestimates of pT181-tau levels in human biofluids and brain samples, depending on the distribution of pT181-tau species in the biospecimen as well as the recognition properties of the antibody. Accordingly, AT270 would be preferred over D9F4G or E4G7X for studies aimed at quantifying *total* pT181-tau species, including variants with or without neighboring phosphorylated residues. Conversely, despite their lower affinity compared to AT270, D9F4G and E4G7X can serve as valuable reagents in assays designed to specifically detect pT181-tau variants without neighboring modifications at S184 and S185. Moreover, comparative analyses using AT270 and either D9F4G or E4G7X could provide an estimate of the extent to which pT181-tau in a given biological sample is also phosphorylated at S184 or S185.

### Cross-Reactivity of AT270 with pT175-Tau

A previous study revealed that AT270 can bind to tau peptides phosphorylated on residue T175 in the absence of pT181, albeit with reduced affinity.^16, 31^ Based on these earlier findings, we assessed the binding specificity of all three pT181-tau antibodies by measuring their affinity for tau peptides singly phosphorylated at neighboring sites on either side of T181, i.e., on residues T175 and S184. Consistent with the earlier report, we found that AT270 bound with lower affinity to the tau peptide phosphorylated on T175 versus T181, with estimated K_D_ values of 248 ± 111 pM and 63 ± 13 pM, respectively (mean ± SEM; Figure S5A, B and Table S1). An increase in the values of both k_on_ and k_off_ was observed for AT270 binding to the pT175-tau versus pT181-tau peptide (Figure S5C, D). However, the magnitude of the increase was markedly greater for k_off_ compared to k_on_ (∼8.6-fold versus ∼2.1-fold, respectively), accounting for the reduced affinity of AT270 for the pT175 target. In contrast to its interaction with the pT175-tau peptide, AT270 exhibited no measurable affinity for the tau peptide singly phosphorylated on residue S184 (data not shown). Moreover, unlike AT270, D9F4G and E4G7X failed to bind to either the pT175-tau or pS184-tau peptide (data not shown). These data indicate that D9F4G and E4G7X have higher specificity for pT181-tau relative to pT175-tau compared to AT270.

The fact that AT270 can bind to pT175 in the absence of pT181 further complicates the interpretation of data from biomarker analyses carried out using this antibody, as it suggests that pT181-tau levels measured in biospecimens containing both pT181- and pT175-tau species using AT270-based assays could in fact be overestimates. Consistent with this idea, MS analyses revealed that tau was phosphorylated on T175 in AD brains, but not in healthy brains,^24^ and that pT175-tau was also present in the CSF of AD patients.^26^ These observations further emphasize the importance of antibody validation in terms of the specificity of binding to tau variants phosphorylated on T175 versus T181 (or at both sites). Moreover, our findings suggest that antibodies with no measurable affinity for pT175-tau (such as D9F4G or E4G7X) can serve as useful tools for discerning whether AT270 immunoreactivity observe in brain specimens or CSF reflects the presence of pT181- or pT175-tau.

### Effects of Neighboring Phosphomimetic Mutations on pT181-Tau Antibody Affinities

Our next objective was to measure the affinities of the pT181-tau antibodies for pT181-tau peptides with a phosphomimetic mutation at position 175, 184, or 185, with the aim of identifying a reliable strategy to test the effects of phosphomimetic substitutions on antibody binding to the full-length, pT181-tau protein. Using our BLI assay, we first measured the affinities of AT270, D9F4G, and E4G7X for pT181-tau peptides with a phosphomimetic aspartate or glutamate substitution at position 175 (T175D/pT181 and T175E/pT181). We found that all three antibodies bound to both pT181-tau peptides with a phosphomimetic mutation at position 175 with affinities that closely resembled those observed during binding to the pT175/pT181-tau peptide (Figure 4 and Table S1). Our observation that the phosphomimetic substitutions at position 175 had no impact on antibody affinity was consistent with the results of T175 phosphorylation.

**Figure 4.**
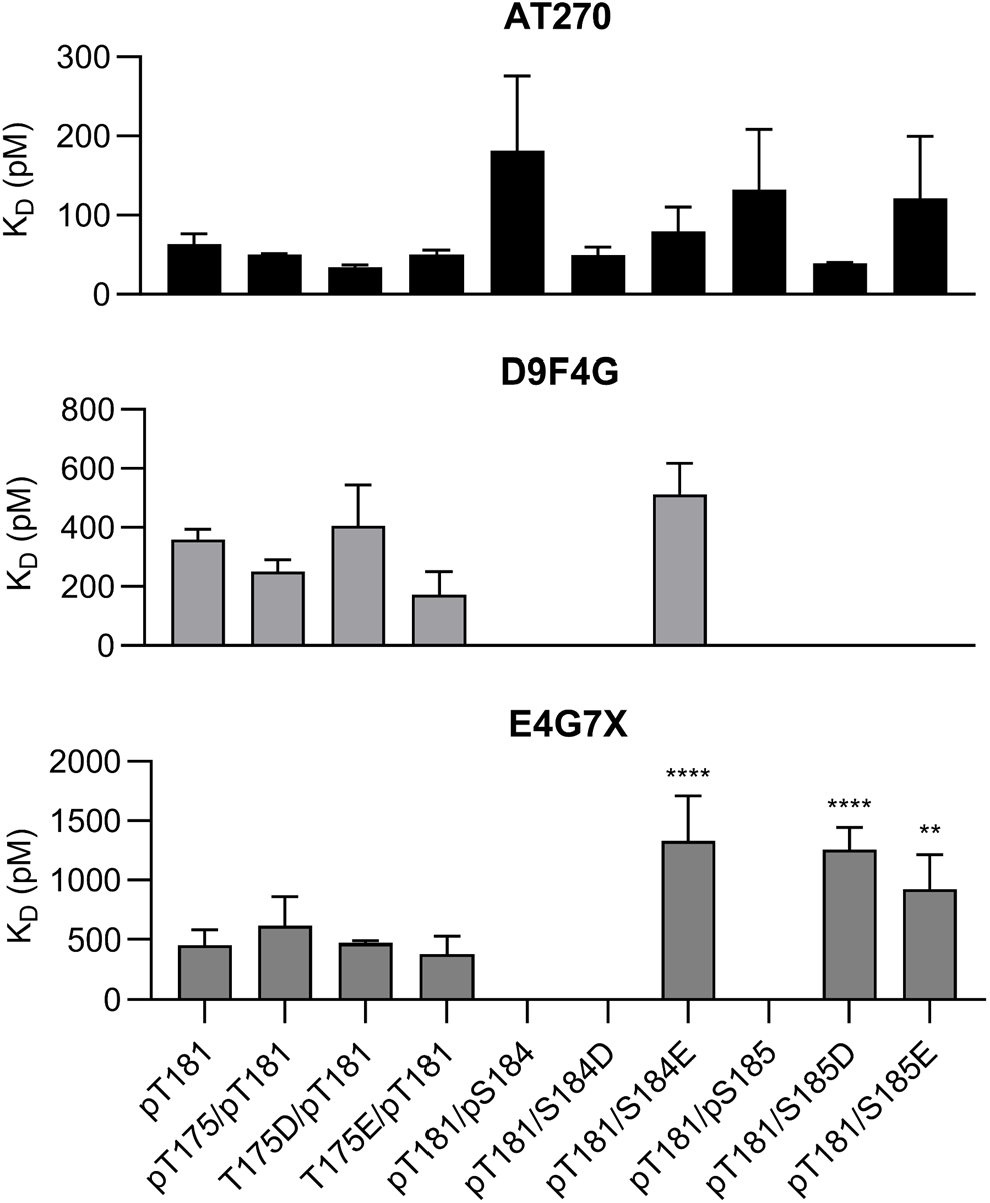
The affinities of pT181-tau antibodies for the anti-genic peptide are affected to different extents by neighboring phosphomimetic mutations. Bar graph showing equilibrium dissociation constants (K_D_ values) determined for AT270, D9F4G, and E4G7X with respect to pT181-tau peptides with a phosphomimetic mutation. Aspartate substitutions at positions 184 and 185 mimicked the effects of phosphorylation at these sites on the binding of AT270, D9F4G, and E4G7X to pT181-tau. Values are expressed as the mean ± SEM (n = 2-13) and are also listed in Table S1; ^**^p ≤ 0.01, ^****^p ≤ 0.0001 *versus* pT181, one-way ANOVA followed by Dunnett’s multiple comparisons *post hoc* test. The absence of a bar indicates that no measurable antibody binding was detected. The data obtained for peptides pT181, pT175/pT181, pT181/pS184, and pT181/pS185 are the same as those shown in Fig. 3.

Next, we measured the affinities of AT270, D9F4G, and E4G7X for pT181-tau peptides with a phosphomimetic aspartate or glutamate substitution at position 184 or 185 (pT181/S184D, pT181/S184E, pT181/S185D, and pT181/S185E). We found that AT270 displayed a similar affinity for all four peptides compared to the unsubstituted pT181-tau peptide, consistent with our observation that AT270 binding to the pT181 target was largely unaffected by S184 or S185 phosphorylation (Figure 4 and Table S1). The presence of aspartate at position 184 or of aspartate or glutamate at position 185 inhibited the binding of D9F4G to the pT181-tau peptide, consistent with this antibody’s inability to bind to both pT181/pS184- and pT181/pS185-tau peptides. An aspartate substitution at position 184 or 185 also eliminated or reduced (respectively) the binding of E4G7X to the pT181-tau peptide, similar to the inhibitory effect of phosphorylation at each site. Conversely, the S184E mutation failed to eliminate the binding of D9F4G or E4G7X to the pT181-tau peptide, and the S185E substitution had no impact on the affinity of E4G7X for the pT181 target.

Collectively, these results indicated that an aspartate substitution at position 184 or 185 mimicked the disruptive effects of phosphorylation at either site on D9F4G or E4G7X binding to the pT181-tau peptide more closely than the corresponding glutamate substitution. Accordingly, the S184D and S185D phosphomimetic mutations were prioritized for subsequent studies aimed at assessing the effects of neighboring phosphorylation events on the affinities of the three antibodies for full-length pT181-tau.

### Effects of Neighboring Phosphomimetic Mutations on the Binding of pT181-Tau Antibodies to the Full-Length Protein Target

A final set of experiments was aimed at determining the effects of neighboring phosphomimetic mutations on the binding of the three pT181-tau antibodies to the target antigen in the context of the full-length (2N4R) protein. In a previous study, Steen and colleagues^32^ found that a 2N4R variant generated using a baculovirus/Sf9 insect cell expression system (referred to here as ‘Sf9-tau’) was extensively phosphorylated on up to 20 residues, including T181 (>50%), but not T175, S184, or S185 (≤2%). Accordingly, we used the same approach to produce 2N4R WT, S184D, and S185D in the pT181 form. We confirmed via LC-MS/MS that all three Sf9-tau variants had a similar phosphorylation profile as that previously reported by Steen and colleagues^32^ as outlined above. Additional LC-MS/MS data revealed that a tau-2N4R variant generated using an *E. coli* expression system was completely unphosphorylated and, therefore, could serve as a negative control for experiments aimed at monitoring the binding of the pT181-tau antibodies to the full-length protein targets.

Next, we determined the relative affinities of each of our three pT181-tau antibodies for Sf9-tau WT, S184D, or S185D via Western blot analysis. Bands of similar intensity corresponding to the pT181 form of all three 2N4R variants were evident on the blot probed with AT270 (Figures 5 and S6), consistent with the similar affinities of AT270 for the corresponding pT181-tau peptides as determined using the BLI assay (Figure 4 and Table S1). In contrast, a band corresponding to pT181-2N4R was only detected in the lane loaded with Sf9-WT (but not -S184D or -S185D) on the blot probed with D9F4G or E4G7X (Figure 5). This result was also in agreement with the relative affinities of the antibodies for the corresponding pT181-tau peptides as determined via BLI, except that the BLI data revealed a measurable amount of E4G7X binding to the pT181/S185D peptide (Figure 4 and Table S1), albeit with a reduced affinity compared to that seen with the unsubstituted pT181 target. Although we observed a lack of E4G7X immunoreactivity to Sf9-tau-S185D on three replicate blots, a fourth blot showed evidence of weak E4G7X binding to this target (Figure S7), consistent with the BLI data. Together, these results suggest that the binding of pT181-tau antibodies to the target antigen is disrupted to different extents by neighboring phosphorylation events in the context of the full-length protein.

**Figure 5.**
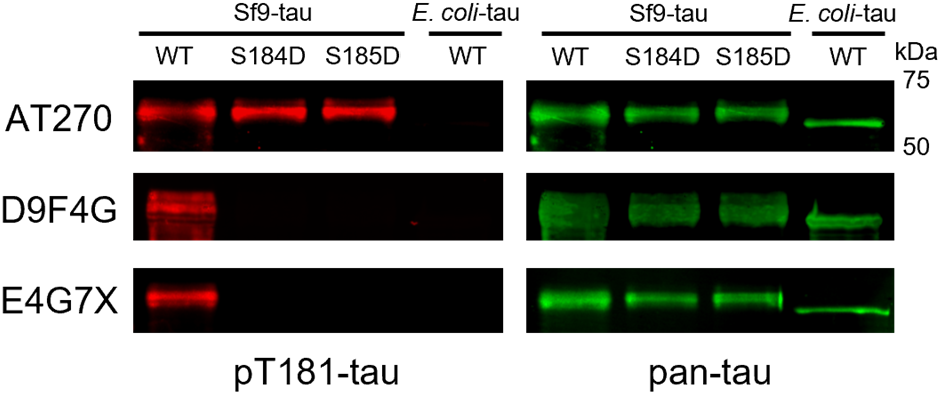
Neighboring phosphomimetic mutations interfere with the binding of D9F4G and E4G7X (but not AT270) to full-length pT181-tau. Recombinant tau variants (WT, S184D, and S185D 2N4R purified from Sf9 cells, and WT 2N4R purified from *E. coli*) were analyzed via Western blotting. Shown are images of Western blots probed with primary antibodies specific for pT181-tau (AT270, D9F4G, or E4G7X; left panel) or a primary antibody specific for total tau (‘pan-tau’; right panel). Images are representative of at least four replicate blots.

### Implications – BLI Assay as a Tool to Validate the Affinity and Specificity of pT181-Tau Antibodies

Our BLI assay offers several advantages over other antibody-based detection methods. In particular, it can be used to determine k_on_, k_off_, and K_D_ values rather than K_D_ values alone, enabling a more detailed understanding of molecular perturbations leading to changes in the affinities of pT181-tau antibodies for their cognate antigen. Details of antibody binding kinetics inferred from BLI measurements can also be used to assess antibody avidity, a quantity inversely related to k_off_ that provides valuable insights into the efficiency of antibody functional responses.^33, 34^ Although ELISA is well-suited to measuring pT181-tau levels in human biospecimens, it cannot be used to monitor the real-time kinetics of antibody-antigen interactions, or to determine whether PTMs have an impact on the association or dissociation phases of anti-body-antigen binding. Additionally, although peptide arrays offer a cost-effective means of screening antibodies against numerous putative targets, the data obtained using array-based methods are semi-quantitative and provide no information about binding kinetics. Similarly, immunoblotting is a semi-quantitative approach that cannot be used to determine K_D_ values or assess the kinetics of antibody-antigen binding. Other advantages of our BLI assay are that it is a label-free method, it affords high sensitivity (enabling the detection of picomolar-level differences in tau antibody affinities), and it can be adapted to a high-throughput format involving low samples volumes (e.g., a volume of only 40 μL is required in each well of a 384-well tilted-bottom plate).

In this study, we demonstrated the utility of our BLI assay in determining the effects of neighboring PTMs on the binding of tau antibodies to their target antigen. Moreover, we showed that our assay could yield valuable insight into the ability of a phosphomimetic mutation at a particular site in the tau protein to mimic the effects of phosphorylation at the same site on antibody binding. In addition to being a validation tool for tau antibody specificity, our BLI assay could potentially be used to measure levels of p-tau variants in biological samples when run in an alternative format involving the immobilization of biotin-conjugated p-tau antibodies to streptavidin biosensors.

By providing information about the antibody binding kinetics, our BLI assay is ideally suited to addressing challenges associated with antibody cross-reactivity. Using our assay, we found that AT270 has significantly higher k_on_ and k_off_ values for the binding of pT175-versus pT181-tau peptide. This observation implies that we could determine whether an AT270-immunoreactive p-tau variant detected in a human biospecimen via BLI consisted of pT175- or pT181-tau by examining the antibody binding kinetics. In contrast, other methods that are limited to reporting target antigen concentrations in the absence of kinetic data (e.g., ELISA^16, 31, 35^) could not be used to address this question with AT270, although such methods could be used with D9F4G or E4G7X given that these antibodies have no detectable affinity for pT175-tau.

## CONCLUSIONS

This study represents a novel application of BLI to determine the effects of neighboring PTMs on the binding of pT181-tau antibodies to their cognate antigen. The BLI assay described herein is a powerful method to characterize antibodies specific for different tau variants in terms of their binding affinities, kinetics, and specificities. Our findings demonstrate that the binding of pT181-tau antibodies to their target is influenced by the phosphorylation of neighboring residues, underscoring the importance of selecting well-characterized antibodies with specific recognition properties for accurate AD biomarker quantitation.

## ASSOCIATED CONTENT

### Supporting Information

The Supporting Information is available free of charge on the ACS Publications website.

Additional experimental procedures and figures are provided in the supporting material file (PDF).

Experimental procedures – LC-MS/MS analysis

Fig. S1. BLI assay design.

Fig. S2. BLI datasets illustrating the approach used to determine optimal pT181-tau peptide loading concentrations.

Fig. S3. BLI datasets illustrating the approach used to determine the optimal range of pT181-tau antibody concentrations.

Fig. S4. BLI datasets showing a lack of measurable binding of pT181-tau antibodies to the unphosphorylated tau peptide.

Fig. S5. Measurement of AT270 affinity for the pT175-tau peptide.

Fig. S6. Neighboring phosphomimetic mutations do not interfere with the binding of AT270 to full-length pT181-tau.

Fig. S7. Western blot image showing evidence of weak E4G7X binding to full-length S185D 2N4R.

Table S1. K_D_ values for the binding of pT181-tau antibodies to phospho-tau target peptides.

## Author Contributions

S.M.: conceptualization, formal analysis, investigation, methodology, visualization, writing – original draft, writing – review & editing. R.M.: formal analysis, investigation, methodology, writing – review & editing. U.K.A.: formal analysis, investigation, methodology, writing – review & editing. T.K.-U.: conceptualization, funding acquisition, resources, writing – review & editing. J.-C.R.: conceptualization, funding acquisition, project administration, resources, supervision, writing – review & editing.

### Notes

The authors declare no competing financial interest.

## ACKNOWLEDGMENTS

This work was supported by grants from Eli Lilly and Company and from the National Science Foundation (1937986-CBET). We thank the MEPEP, Proteomics, and Chemical Genomics facilities at Purdue University for technical support with protein purification, MS analyses, and BLI analyses, respectively. We also thank Thorsten Wiederhold and Arica Aiello for providing the E4G7X antibody, and David Eliezer for providing the pRK172-h-tau-2N4R bacterial expression plasmid.

Authors are required to submit a graphic entry for the Table of Contents (TOC) that, in conjunction with the manuscript title, should give the reader a representative idea of one of the following: A key structure, reaction, equation, concept, or theorem, etc., that is discussed in the manuscript. Consult the journal’s Instructions for Authors for TOC graphic specifications.

## Insert Table of Contents artwork here

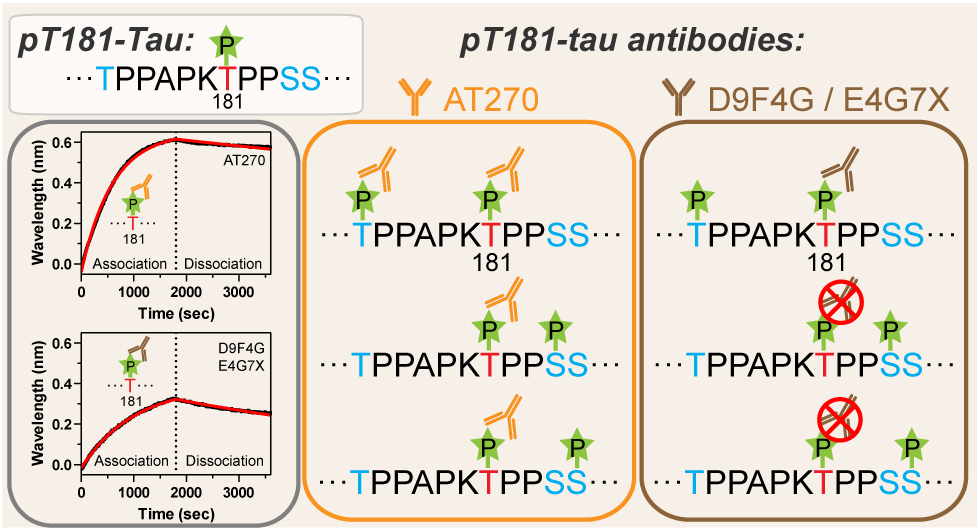

