## Supporting Information for "Effects of Neighboring Phosphorylation Events on the Affinities of pT181-Tau Antibodies"

### I. Experimental procedures

#### Sample preparation for MS analysis

Protein samples (5 µg) were reconstituted in 50 µL of an 8 M urea solution prepared in 50 mM ammonium bicarbonate (ABC) and incubated at 25 °C for 20 min. DTT was then added to a final concentration of 5 mM, and samples were incubated at 37 °C in a thermomixer for 30 min. For cysteine alkylation, iodoacetamide (IAA) was added to a final concentration of 20 mM, and samples were vortexed at 25 °C, in the dark, for 45 min. The remaining IAA was then quenched with excess DTT for 10 min at 25 °C. Trypsin (0.5 µg) was added to each sample, and the total volume was adjusted to 400 µL with 5% (v/v) acetonitrile (ACN) in 50 mM ABC. Protein digestion was carried out at 37 °C overnight, and peptide cleanup was carried out with peptide desalting columns. Samples were then dried and reconstituted in 3% (v/v) ACN and 0.1% (v/v) Formic Acid (FA) in $H_2O$ for MS analysis.

#### Mass spectrometry

MRM analyses were performed with a TSQ Endura instrument, coupled with an Ultimate 3000 HPLC. Reverse-phase separation was performed with an Aurora Ultimate analytical column (25 cm), packed with 1.7 µm 120 Å C18 medium (IonOptics), using a 60-min gradient. Peptides were first loaded into the column with 98% Mobile phase A (0.1% v/v FA in $H_2O$) and 2% mobile phase B (80% v/v ACN with 0.1% v/v FA in $H_2O$). At minute 2.6, mobile phase B was increased to 8%. Mobile phase B then was linearly increased to 27% for 37.4 min, followed by an increase to 45% B after 5 min. Mobile phase B was subsequentially increased to 100% at minute 50, and held constant for a total of 5 min. Peptide transitions were generated using Skyline, accounting for all possible phospho-STY modification sites. Up to 1 miscleavage was allowed per peptide. Q1 resolution (FWHM) was set to 0.7, and Q3 resolution was set to 1.2. Optimal collision energies predicted by Skyline were used for each peptide analyzed. Raw data were then imported and analyzed within the Skyline platform.

#### II. Supporting Data

| Step | Time (sec) | Microplate column number |
|---|---|---|
| Equilibrium | 60 | 1 |
| Loading | 300 | 3 |
| Baseline | 300 | 5 |
| Baseline | 10 | 7 |
| Association | 1800 | 9 |
| Dissociation | 1800 | 7 |

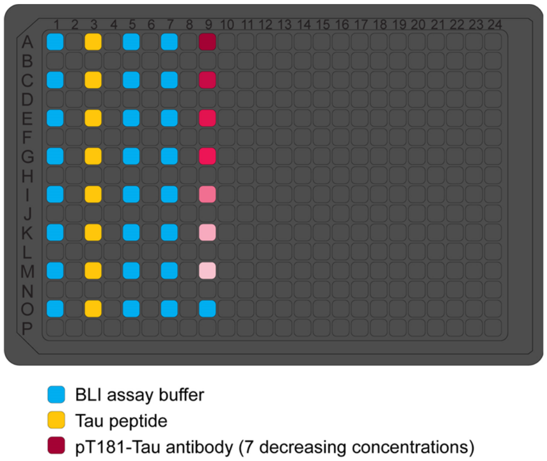


**Figure S1. BLI assay design.** (Left) Table listing the different steps of the assay, as well as the incubation time and microplate column number for each step. (Right) Illustration of the 384-well plate setup used for the assay. Each well highlighted in color contained 40 µL of BLI assay buffer. A subset of wells was supplemented with tau peptide (final concentration,10 ng/mL; column 3) or pT181-tau antibody (diluted to one of seven concentrations; column 9). The background signal was determined by measuring the wavelength shift of a biosensor incubated with tau peptide in the absence of antibody.

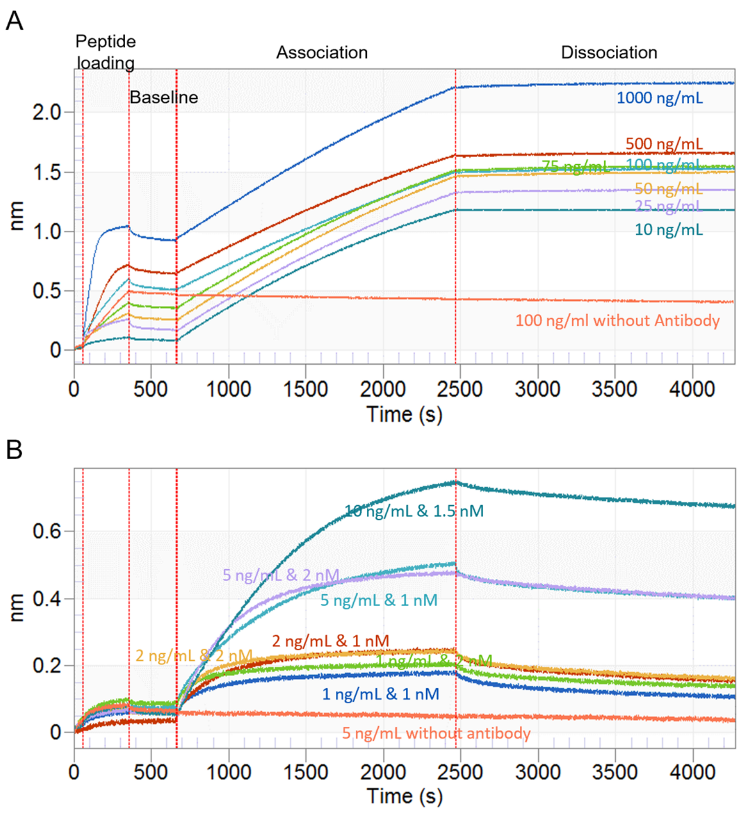


**Figure S2. BLI datasets illustrating the approach used to determine optimal pT181-tau peptide loading concentrations.** (A) Sensorgram traces obtained for biosensors loaded with pT181-tau peptide (10 to 1000 ng/mL) and incubated with the pT181-tau antibody AT270 (1.5 nM) (association phase) followed by assay buffer without antibody (dissociation phase). (B) Sensorgram traces obtained for biosensors loaded with pT181-tau peptide (1 to 10 ng/mL) and incubated with AT270 antibody at concentrations of 1, 1.5, and 2 nM, followed by assay buffer without antibody.

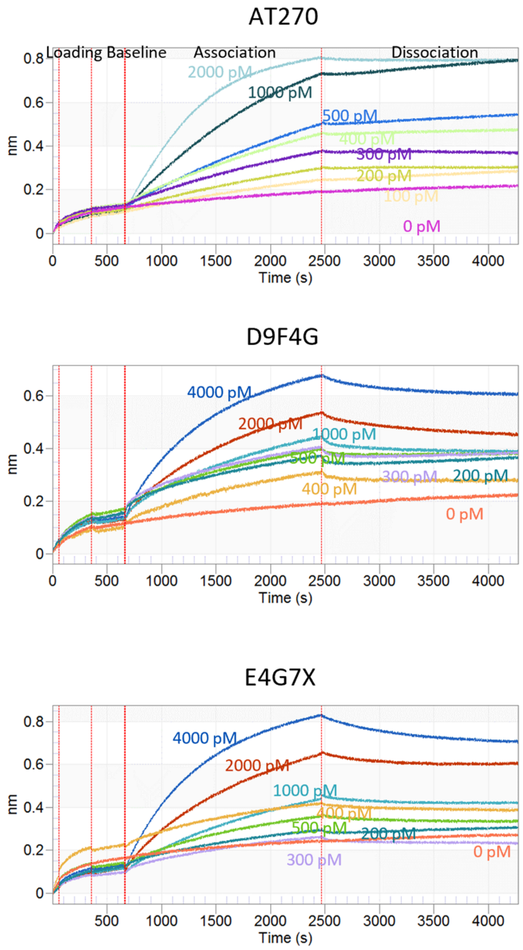


**Figure S3. BLI datasets illustrating the approach used to determine the optimal range of pT181-tau antibody concentrations.** Sensorgram traces obtained for biosensors loaded with pT181-tau peptide (10 ng/mL) and incubated with AT270 (top), D9F4G (middle), or E4G7X (bottom) at the indicated concentrations (association phase), followed by assay buffer without antibody (dissociation phase).

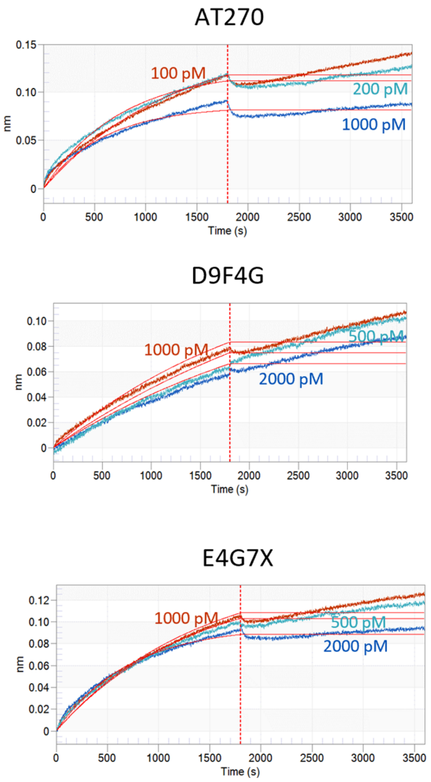


**Figure S4. BLI datasets showing a lack of measurable binding of pT181-tau antibodies to the unphosphorylated tau peptide.** The sensorgrams include raw-data traces and best-fit curves (red) corresponding to the association and dissociation of AT270 (top), D9F4G (middle), or E4G7X (bottom) at different concentrations.

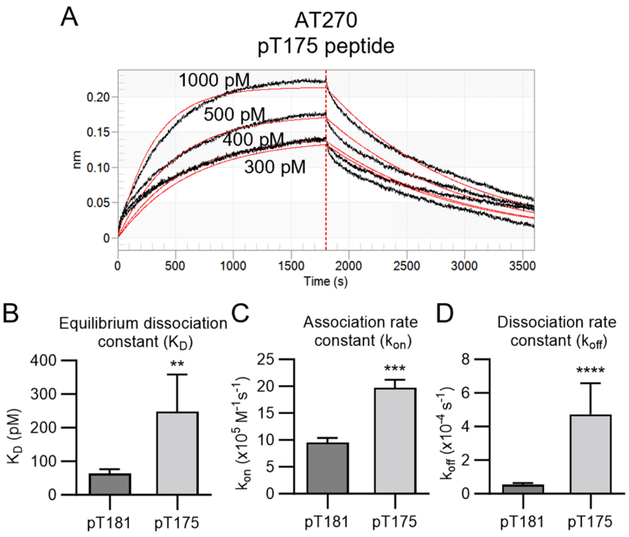


**Figure S5. Measurement of AT270 affinity for the pT175-tau peptide.** (A) Sensorgram showing raw-data traces (black) and best-fit curves (red) corresponding to the association and dissociation of AT270 upon incubation at different concentrations with biosensors loaded with pT175-tau peptide. (B-D) Bar graphs showing equilibrium dissociation constants ($K_D$ values) (B), association rate constants ($k_{on}$ values) (C), and dissociation rate constants ($k_{off}$ values) (D) determined for AT270 with respect to the pT181- and pT175-tau peptides. AT270 had a measurable affinity for the pT175-tau peptide, despite the lack of a phosphate group on residue T181. The measured $K_D$, $k_{on}$, and $k_{off}$ values were greater for the binding of AT270 to the pT175-tau peptide compared to the pT181-tau peptide. Values are expressed as the mean ± SEM (n = 2-12); $^{**}p \leq 0.01$, $^{***}p \leq 0.001$, $^{****}p \leq 0.0001$, unpaired two-tailed t-test. The data obtained for peptide pT181 are the same as those shown in Fig. 2.

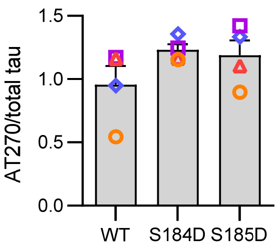


**Figure S6. Neighboring phosphomimetic mutations do not interfere with the binding of AT270 to full-length pT181-tau.** Recombinant tau variants (WT, S184D, and S185D 2N4R purified from Sf9 cells) were analyzed via Western blotting using AT270 or pan-tau as the primary antibody (see Figure 5 of the main text). The bar graph shows the ratio of the AT270 to pan-tau band intensities determined for each variant via densitometric analysis of the blot. Values are expressed as the mean ± SEM (n = 4). Symbols of the same shape and color correspond to data obtained from the same blot.

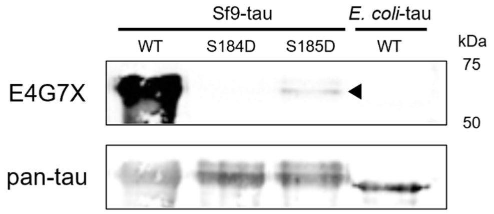


**Figure S7. Western blot image showing evidence of weak E4G7X binding to full-length S185D 2N4R.** Recombinant tau variants (WT, S184D, and S185D 2N4R purified from Sf9 cells, and WT 2N4R purified from *E. coli*) were analyzed via Western blotting using E4G7X or pan-tau as the primary antibody (see Figure 5 of the main text). This was the only blot out of 4 replicates where we detected weak E4G7X binding to Sf9-tau-S185D.

**Table S1. $K_D$ values for the binding of pT181-tau antibodies to phospho-tau target peptides**

| Peptide | $K_D$ (pM)[1] | | |
|---|---|---|---|
| | AT270 | D9F4G | E4G7X |
| pT181 | 63 ± 13 | 360 ± 35 | 459 ± 37 |
| pT175/pT181 | 50 ± 1 | 252 ± 39 | 614 ± 175 |
| pT181/pS184 | 182 ± 94 | nd | nd |
| pT181/pS185 | 132 ± 77 | nd | nd |
| pT175/pT181/pS184 | 103 ± 9 | nd | nd |
| pT175/pT181/pS185 | 80 ± 12 | nd | nd |
| pT181/pS184/pS185 | 107 ± 32 | nd | nd |
| T175D/pT181 | 34 ± 3 | 405 ± 139 | 474 ± 12 |
| T175E/pT181 | 50 ± 6 | 173 ± 77 | 386 ± 100 |
| pT181/S184D | 50 ± 10 | nd | nd |
| pT181/S184E | 80 ± 30 | 513 ± 106 | 1330 ± 270 |
| pT181/S185D | 39 ± 2 | nd | 1257 ± 107 |
| pT181/S185E | 121 ± 79 | nd | 921 ± 146 |
| pT175 | 248 ± 111 | nd | nd |

[1]Values from Figures 3 and 4 are expressed as the mean ± SEM (n = 2-13); nd, antibody binding not detected.
